# Intercellular signaling reinforces single-cell level phenotypic transitions and facilitates robust re-equilibrium of heterogeneous cancer cell populations

**DOI:** 10.1101/2025.01.03.631250

**Authors:** Daniel Lopez, Darren R Tyson, Tian Hong

## Abstract

**Background:** Cancer cells within tumors exhibit a wide range of phenotypic states driven by non-genetic mechanisms in addition to extensively studied genetic alterations. Conversions among cancer cell states can result in intratumoral heterogeneity which contributes to metastasis and development of drug resistance. However, mechanisms underlying the initiation and/or maintenance of such phenotypic plasticity are poorly understood. In particular, the role of intercellular communications in phenotypic plasticity remains elusive.

**Methods:** In this study, we employ a multiscale inference-based approach using single-cell RNA sequencing (scRNA-seq) data to explore how intercellular interactions influence phenotypic dynamics of cancer cells, particularly cancers undergoing epithelial-mesenchymal transition. In addition, we use mathematical models based on our data-driven findings to interrogate the roles of intercellular communications at the cell populations from the viewpoint of dynamical systems.

**Results:** Our inference approach reveals that signaling interactions between cancerous cells in small cell lung cancer (SCLC) result in the reinforcement of the phenotypic transition in single cells and the maintenance of population-level intratumoral heterogeneity. Additionally, we find a recurring signaling pattern across multiple types of cancer in which the mesenchymal-like subtypes utilize signals from other subtypes to support its new phenotype, further promoting the intratumoral heterogeneity. Our models show that inter-subtype communication both accelerates the development of heterogeneous tumor populations and confers robustness to their steady state phenotypic compositions.

**Conclusions:** Our work highlights the critical role of intercellular signaling in sustaining intratumoral heterogeneity, and our approach of computational analysis of scRNA-seq data can infer inter- and intra-cellular signaling networks in a holistic manner.

## Background

Most tumors develop and evolve as complex ecosystems under strong environmental selective pressures, leading to a unique collection of cancer cells that exhibit a wide range of genotypic and phenotypic characteristics^1,2^. This intratumoral heterogeneity promotes aggressive disease progression, increased resistance to therapeutic interventions, and poor overall survival^1,3–5^. While genetic diversity is a well-known driver of intratumoral heterogeneity^6^, there is increasing evidence that non-genetic mechanisms, such as epigenetic, transcriptional, and/or translational changes, also significantly contribute to the intratumoral heterogeneity and disease progression^4,5,7,8^. These non-genetic mechanisms can create distinct cancer cell states through a process called phenotypic plasticity, where cells are dynamic, reversible, and responsive to regulatory changes^3,9,10^.

Phenotypic plasticity has recently been recognized as a hallmark of cancer and a key driver of tumor aggressiveness^3^. It influences various cellular behaviors in cancer, including stemness and differentiation, drug-sensitive and drug-resistant states, and transitions between epithelial and mesenchymal cell-states^11^. Increasing efforts are being made to characterize the intrinsic cellular factors that drive phenotypic plasticity^12–14^. However, despite extensive molecular characterization, the dynamics of phenotypic plasticity at the single-cell and population levels remain largely unclear. This is particularly true regarding how non-cell-autonomous effects regulate intratumoral heterogeneity and whether the intercellular communication between different cell states stabilizes or destabilizes these phenotypes.

One such cancer where phenotypic plasticity is particularly evident is small cell lung cancer (SCLC)^15,16^. SCLC is a neuroendocrine (NE) carcinoma that constitutes approximately 15% of all lung cancer cases and has a dismal five-year survival rate of less than 7%^17^. Despite the high similarity to pulmonary NE cells and having highly consistent morphological characteristics, SCLC presents substantial inter- and intratumoral heterogeneity, featuring distinct molecular subtypes with varied biological behaviors^18–22^. These subtypes are categorized based on the enriched expression of one of four transcription factors (TFs): *ASCL1*, *NEUROD1*, *POU2F3*, *YAP1*^23^. Furthermore, these subtypes delineate into two overarching categories, NE (SCLC-A2, -A, and -N) and non-NE (SCLC-P and -Y), with the NE subtypes typically exhibiting some level of ASCL1 expression while non-NE counterparts do not. Recent studies have demonstrated that SCLC tumors will often comprise multiple cell types, with the different subtypes cooperating to drive tumorgenicity^21,22^. The dynamic regulation of TFs regulates intratumoral compositions, and this diversity is essential as different subtypes play distinct biological roles, impacting therapeutic response^21,22^.

Further highlighting the phenotypic plasticity evident within SCLC, previous work by us^24^ and others^25,26^ has linked the different SCLC subtypes to the epithelial-mesenchymal transition (EMT) program, a cellular process in which cell-cell interactions are remodeled, resulting in cells losing their epithelial properties and assuming a more mesenchymal phenotype.^27^. Within SCLC, the NE subtype SCLC-A2 demonstrates a strong epithelial-like phenotype whereas the other NE subtypes, SCLC-A and SCLC-N, display a partial EMT state (Figure 1A). non-NE subtypes, SCLC-P and SCLC-Y, also display a partial EMT state, albeit with mesenchymal gene expression signatures that differ from that of the NE subtypes. This correspondence with EMT further demonstrates the plastic nature of this cancer.

**Figure 1.**
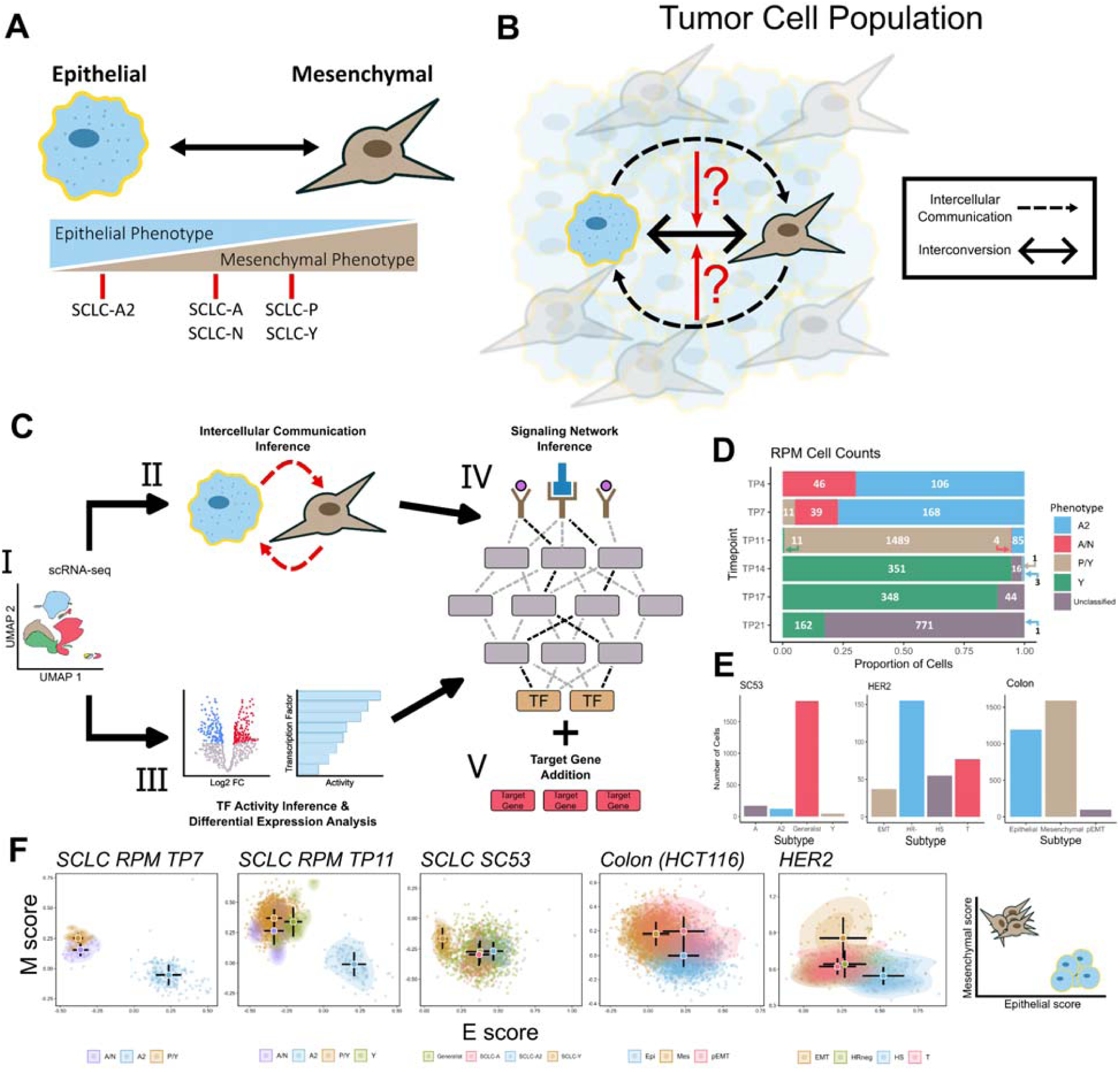
Multiscale Inference Approach to Investigate Role of Intercellular Communication on Cellular Plasticity. A) Schematic depicting EMT and correspondence to SCLC subtypes. Subtypes are characterized based on transcriptomic expression. B) Investigating whether intercellular communications affect the cell state transitions between subtypes and the overall intratumoral heterogeneity. Black dashed arrows represent inferred cell-cell communication and the black, double-arrow represents the interconversion between subtypes. C) I: Pre-processed and annotated scRNA-seq data. Pipeline is applied to four different datasets across three different cancers II: Inference of cell-cell communications using CellChat. III: Differential expression and transcription factor activity analysis. IV: Signaling network inference via CORNETO is performed using th receptor probability and transcription factor activity scores as inputs. V: Differentially expressed SCLC and EMT gene markers are added to the network based on filtering criteria. D) Filled bar chart showing the cell counts for the SCLC RPM dataset at TP7. X-axis represents the proportion of cells and y-axis is the time point. The cell numbers for each cell type are shown. E) Cell counts for SCLC SC53, HER2 and colon cancer datasets, respectively. The cell type annotations are from the original authors. F) Scatter plots of single-sample gene set enrichment analysis (ssGSEA). Each dot represents a cell in the dataset. X- and y-axis represent enrichment scores of epithelial and mesenchymal genes, respectively. Summary of the results depicting the arrangement of epithelial and mesenchymal cells on the far right.

Most research on SCLC has focused on describing its differing cell states, with very few studies attempting to define these states at the single-cell level. Consequently, there is still much to learn about the transitions between different cell states and whether intercellular communication influences the intratumoral heterogeneity. Given the plasticity present within SCLC, we utilized single-cell RNA-sequencing (scRNA-seq) SCLC data to investigate whether intercellular communications influence cell-fate transitions, and if so, whether these extracellular signals reinforce the current phenotype of a cell or push it towards a different phenotype.

To investigate whether intercellular communications play a role in driving intratumoral heterogeneity within SCLC, we adapted a single-cell multiscale inference-based approach that integrates both intercellular interactions and intracellular signaling dynamics. By linking inter- and intracellular signaling, we aim to determine if intercellular interactions impact downstream intracellular mechanisms. If cell–cell communication affects downstream signaling, this approach will also reveal whether these interactions drive the cell towards a different phenotype or reinforce its current one (Figure 1B). To explore the effect of cell–cell communications on phenotypic heterogeneity, we utilized scRNA-seq data from *ex vivo* cultured cells obtained from a genetically engineered mouse model that incorporates a constitutively active form of *MYC*, coincident with deletions of *Rb* and *p53* (RPM)^28^. The cells undergo a transition over time in culture from a NE to non-NE state and these time series data enable the exploration of intercellular communications between the NE and non-NE populations at individual time points. Through this approach, we identified the activation of several well-established EMT pathways, revealing a consistent pattern of convergence towards activating mesenchymal genes within the mesenchymal phenotype. Additionally, our analysis revealed that the epithelial phenotype employs both paracrine and autocrine signaling mechanisms to maintain its epithelial state. To see if these mechanisms are consistent across epithelial cancers that involve EMT, we applied this method to colon and breast cancer datasets. The convergence of EMT pathways towards the mesenchymal phenotype is present across all 3 cancers. However, SCLC appears to be unique in its utilization of both paracrine and autocrine signals to sustain its phenotypic state. Overall, our results show recurring roles of intercellular communications in maintaining newly formed cell states, and they shed light into non-cell-autonomous mechanisms of intratumoral heterogeneity.

## Results

### A multiscale inference approach to explore the signaling mechanisms maintaining phenotypic heterogeneity in cancer cell populations

To assess whether cell-cell communications contribute to intratumoral heterogeneity and phenotypic plasticity it is essential to connect intercellular signaling with downstream intracellular processes and determine their overall impact on maintaining phenotypic diversity. To link intercellular communication with intracellular signaling, we adapted a methodology that integrates various approaches to explore both intercellular and intracellular signaling events (see Methods, Figure 1C, and reference^29^). Briefly, we infer the active signaling pathways and ligand–receptor (L–R) interactions to assess intracellular signaling from processed scRNA-seq data^30^. We opted for this approach due to its capacity to incorporate heteromeric complexes and its robustness to noise^31^.

The inferred active signaling pathways are then used to predict transcriptional activity and identify the relative strength of active signaling pathways in each cell state. We then used an integer linear programming tool for contextualizing causal networks^32^ to integrate the transcription factor (TF) activity inference scores, differential expression analysis, and L–R interactions. This approach finds the smallest-sign consistent network that explains the measured inputs and outputs, connecting the receptors to the downstream TFs. To aid in assessing the relationships between EMT states cellular differentiation, we incorporate downstream SCLC^21^ and EMT^33^ target genes in the network—adding specific target genes based on the congruence of the inferred regulatory modes between TF and target gene (i.e. activation or inhibition) and experimental values of gene expression for the target gene. This multiscale integration allows us to capture dynamic changes in signaling networks across different cell states and enables a detailed exploration of the potential signaling mechanisms involved in maintaining the phenotypic heterogeneity within the cancerous population.

We applied this approach to the SCLC RPM dataset, focusing primarily on timepoints 7 and 11 (TP7 & TP11), when both NE and non-NE subtypes are present in relatively high abundance (Figure 1D). Within this dataset, five subtypes were determined via archetype analysis (SCLC-A2, -A/N, -P/Y, -Y and Unclassified), with the SCLC-A2 subtype displaying more epithelial-like properties, while the SCLC-P/Y and SCLC-Y subtypes exhibited more mesenchymal like features, as expected. Unclassified cells are cells which could not be assigned to any of the defined SCLC-A, -N, -P, -Y archetypes (see Methods). We then extended this approach to three additional datasets involving cancers undergoing EMT: SC53 (Human SCLC circulating tumor cells-derived xenograft sample^25^), HCT116 colon cancer cell line^34^, and a HER2 Crainbow mouse^35^(Figure 1E & Supplementary Figure 1). The cell type classifications in these datasets were determined by the authors who generated the data. In SC53 there are four identified subtypes: SCLC-A, -A2, -Y and Generalists. Notably, the SCLC-Y subtype exhibited gene expression profiles consistent with a more mesenchymal-like state. In the colon cancer dataset, three EMT-associated states were characterized: epithelial (Epi), mesenchymal (Mes), and partial-EMT (pEMT). The HER2 dataset has four cell-states that were inferred through trajectory analysis: hormone-sensitive (HS), hormone-receptor negative (HS-), EMT and a transitional (T) state. Given the different strategies of cell type annotation across the datasets, we assessed whether each dataset contained distinguishable epithelial- and mesenchymal-like populations across the different datasets. Through gene set enrichment analysis methods, we revealed a clear separation of epithelial and mesenchymal populations across the different datasets (Figure 1F and Supplementary Figure 2). Additionally, to account for the large number of Unclassified cells in the SCLC RPM dataset, we also examined how adjusting the archetype assignment threshold impacted the clustering results. We found that incorporating more of these cells into the defined subtypes did not significantly alter the overall clustering patterns (Supplementary Figure 3A & B).

### Diverse signaling pathways converge on activating key mesenchymal pathways/genes

We first analyzed the RPM dataset and identified 48 active signaling pathways across the six different timepoints (Supplementary Table 1). Many of these pathways are well-established contributors to EMT in cancer^27,36–57^ (Figure 2A). Notably, we observed a recurring pattern among these EMT pathways wherein they converge towards the mesenchymal non-NE cell types (SCLC-P/Y in Figure 2B). This convergence is mediated by paracrine (NOTCH) signaling from NE to non-NE cells and autocrine signaling within the non-NE cell population (WNT and SPP1).

**Figure 2.**
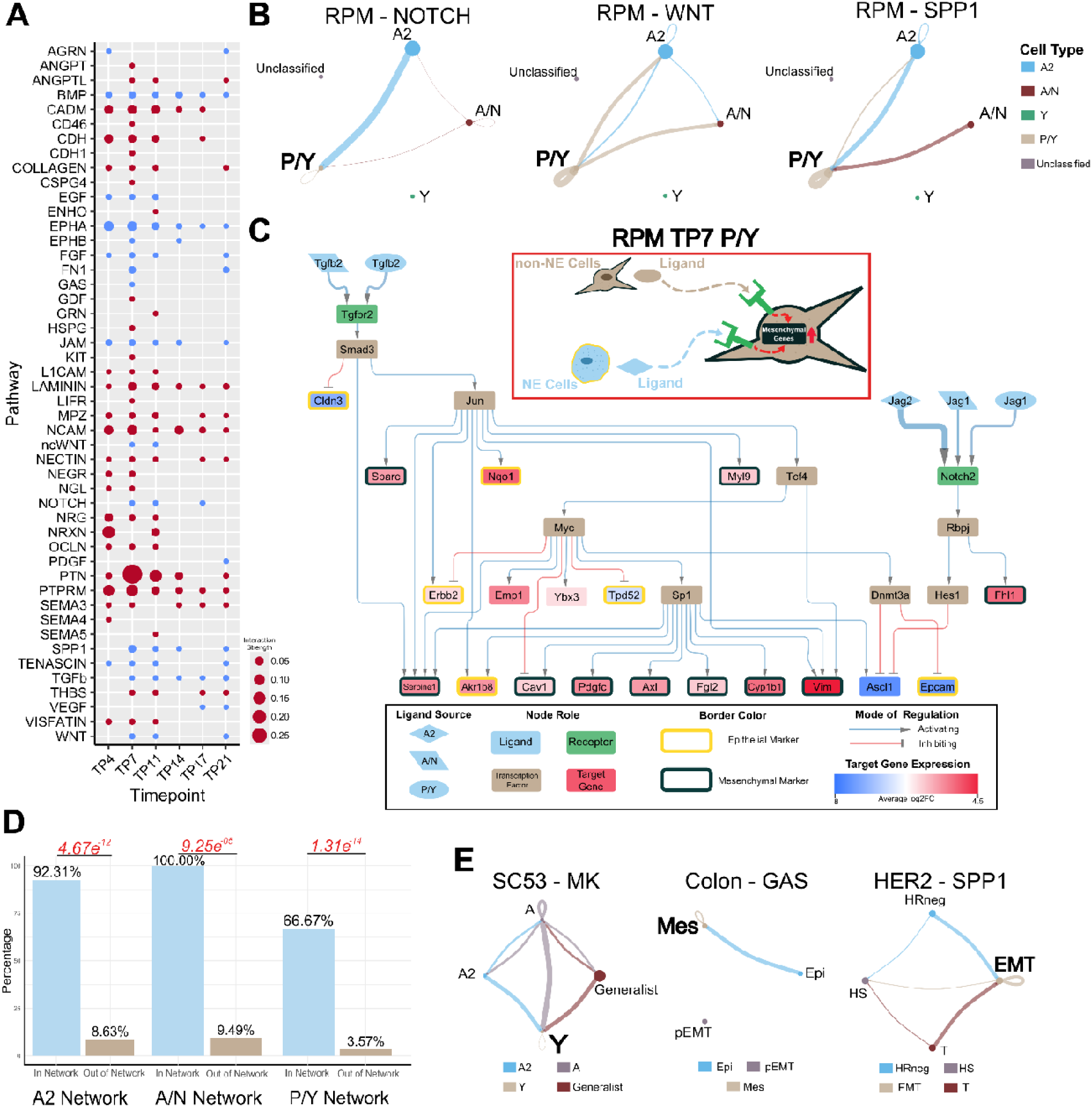
Mesenchymal Phenotypes Utilize Autocrine and Paracrine Signaling to Reinforce SCLC Subtypes. A) Inferred active pathways in the SCLC RPM dataset. Dot size is representative of the relative interaction strength for a given time point. Blue circles are pathways known to be involved in EMT signaling. B) Inferred *CellChat* signaling of pathways known to be involved in EMT at TP7. Dot size is proportional to cell number. The legend on the right contains the cell type each dot represents. Line color represents the source of signal/ligand. Line width represents interaction strength. C) RPM TP7 SCLC-P/ cell type inferred signaling network connecting both intercellular and intracellular signaling. The red box contains an illustrative summary of the network. The black box is a legend for the network. Node color represents the role in the signaling pathway. The shape of the ligand nodes represents the subtype source of the ligand. The edges connecting a ligand to a receptor vary in width corresponding to interaction strength. Average log2 fold-change of target gene expression is colored as indicated by the scale bar. Epithelial target genes are indicated by a yellow border around the node and mesenchymal genes have a black border. D) Proportion of lineage supporting genes included in (In Network) and excluded from (Out of Network) the network. Fisher’s exact test was used to calculate significance. P-values are labeled in red. Odds ratios from left to right: 126.99, inf, 54.08. E) Inferred *CellChat* signaling of EMT pathways from other datasets. Left: Human SCLC CDX. Subtypes are characterized through archetype analysis. Middle: HCT116 colon cancer. Subtype labels are Epi (Epithelial), Mes (Mesenchymal), and pEMT (Partial EMT) and they are from the original publication. Left: HER2 breast cancer mouse. Subtype labels are HS (Hormone-sensitive), HR-negative (Hormone-receptor negative), T (Transitional state), and EMT. The more mesenchymal states in these datasets are, Y, Mes, and EMT, respectively.

To elucidate how the intercellular communication affects intracellular signaling, we constructed a signaling network for the Day 7 SCLC-P/Y cell type that has TGFβ and NOTCH as primary ligands and captured key lineage-supporting target genes (Figure 2C). Within the network, we observed the activation of numerous mesenchymal markers (like vimentin, PDGF-C and Axl) and the inhibition of epithelial markers, especially Epcam. Consistent with previous experimental findings^16^, we noted that the activation of *Myc* and Notch signaling promotes the non-NE SCLC fate. Our network further highlights that the inhibition of *Ascl1*—a NE and SCLC-A & -A2 subtype marker downstream of the *Notch2* receptor—and the activation of *Myc* facilitates the upregulation of mesenchymal markers. Notably, the ligand–receptor interactions captured in the network are well known EMT modulators. Since the ligands involved in the activation of mesenchymal markers within the SCLC-P/Y network originate from SCLC-A2, -A/N and -P/Y cells, our network underscores the contributions of both NE and non-NE cells toward mesenchymal transitions. This approach yielded similar results in the TP11 SCLC-P/Y cell type by capturing the activation of lineage-supporting genes, with both NE and non-NE cells playing a role in activating those genes (Supplementary Figure 4). Additionally, to further assess the robustness of our network results, we decreased the assignment threshold in the archetype analysis to assign more Unclassified cells into one of the SCLC subtype archetypes. While this increased the total network size for the SCLC-P/Y group at TP7, the expanded network still captured a majority of the same nodes as the original network. This indicates that the inclusion of more Unclassified cells does not substantially alter the overall network structure (Supplementary Figure 5).

We then assessed the importance of lineage supporting genes captured in the networks by creating a *2×2* contingency table and determining the ratios of lineage supporting genes included in the inferred signaling networks versus those excluded from the network. We found a higher proportion of lineage supporting genes within the network compared to outside the network (Figure 2D) (Supplementary Table 2), suggesting that the inferred intercellular communication driving the mesenchymal (M) state transcriptional program was not simply due to the random selection of broadly upregulated M genes.

To assess the generalizability of these findings across other cancers undergoing EMT, we performed a similar analysis on three additional scRNA-seq datasets. We observed the activation of intercellular communication-driven mesenchymal pathways converging towards a more mesenchymal phenotype in all three datasets (Figure 2E). However, application of the signaling network pipeline to the colon and HER2 datasets yielded less comprehensive networks compared to those observed in SCLC (Supplementary Figure 6). This discrepancy suggests potential differences in the sensitivity to detect extracellular signals or a reduced influence of the tumor microenvironment in driving phenotypic heterogeneity within these datasets. Overall, our findings demonstrate that the intercellular communications between cancer cells can have varying degrees of influence in maintaining phenotypic diversity. Within SCLC, the intercellular communications between NE and non-NE phenotypes reinforce the non-NE/mesenchymal (SCLC-P/Y or -Y) state.

### A mathematical model predicts the roles of inter-subtype feedback in stabilizing phenotypic compositions of heterogeneous tumor cell populations

Analysis of previously published data showed that the co-existence of epithelial-(E-)like cells and mesenchymal-(M-)like cancer cell subpopulations is a recurring pattern in multiple cancer types (Figure 1F). Through our inference pipeline in this work, we also found a type of intercellular communication shared by multiple cancer types and datasets: cells at the state enriched with M genes receive signals from cells at a more E-like state, and the signals were used to maintain the recently acquired M state (Figure 2). Previous work has shown the functions of autocrine signaling in maintaining cell states^58,59^, but it is unclear how the inter-subtype signals that we found in multiple contexts can influence intratumoral population dynamics. We therefore built a simple mathematical model based on the following dimensionless ordinary differential equations (ODEs)

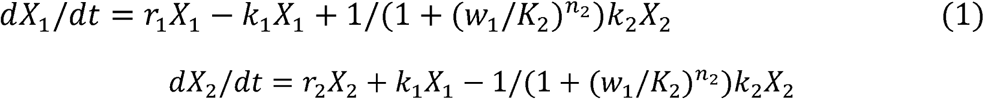

which depict two subpopulations (X_1_ and X_2_) with proliferation rate constants *r*_1_ and *r*_2_ respectively, and basal interconversion rate constants (*k*_1_ and *k*_2_ for X_1_-to-X_2_ conversion and X_2_-to-X_1_ conversion, respectively) (Figure 3A). The signal that originates from X_1_ and received by X_2_ is modeled by an inhibitory Hill function that influences the overall conversion rate from X_2_ to X_1_, as the inferred inter- and intra-cellular activities common to all cancer types. Parameter *K*_2_ determines the threshold of the inactivation and *n*_2_ determines the nonlinearity of the response. *w*_1_ is the abundance of X_1_ relative to the total size of the population (i.e. *w*_1_ = *X*_1_/(*X*_1_ + *X*_2_). This Hill function effectively serves as a feedback mechanism that may influence the dynamics of the cancer cell population. Simply, the model describes the dynamics of a cell population with two discrete states (of any type) with interactions between them.

**Figure 3.**
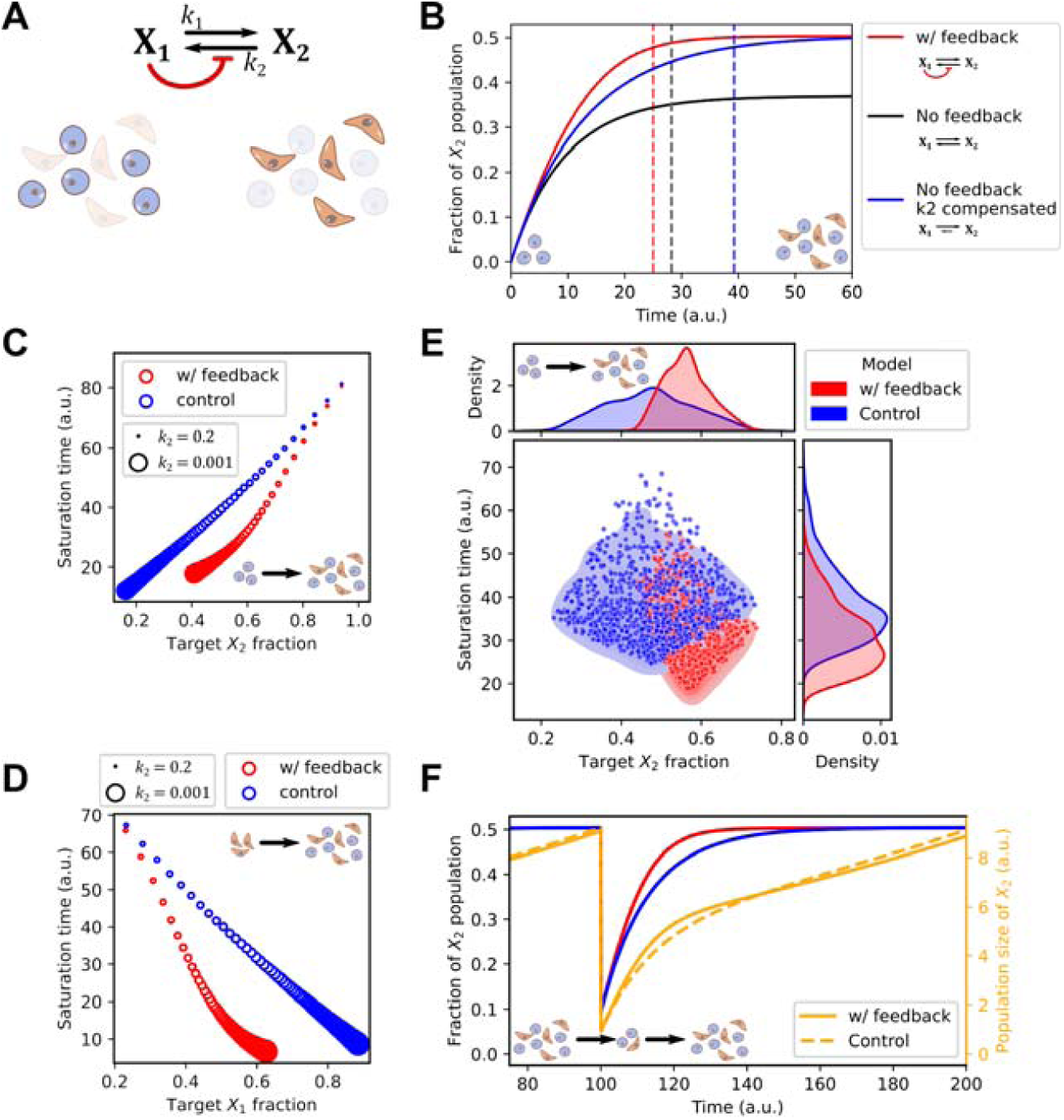
A mathematical model for inter-subtype communication. A) Network diagram and illustration of two cell types (Epithelial-like and Mesenchymal-like). X_1_ and X_2_ are two state variables representing the sizes of the two cell populations, respectively. Black arrows show transitions. Red arrow shows the intercellular communication responsible for maintaining the M-like cell state. B) Time course trajectories (solid curves) for the main model and two perturbed models. Y-axis shows the quantity X_2_/(X_1_+X_2_). Dashed vertical lines show the positions of saturation times, defined as the time at which 95% of the steady state level of the fraction is reached. Initial conditions: X_1_ =10, and X_2_=0. See Methods for parameter values. C) Saturation time as a function of steady state fraction of X_2_. Color code is the same as Panel B. 100 evenly spaced k_2_ values in the indicated range were chosen to achieve various steady state X_2_ fractions for each model. Circle sizes indicate k_2_ values. Initial conditions are parameter values are the same as B. D) Same simulations as in C except for initial conditions. Here, X_1_ =0, and X_2_=10. E) Performance of two models with 1000 sets of randomly chosen values for all parameters with 50% decrease/increase from the basal values as in B. F) Time course trajectories of a fractional killing scenario in which the fractions of the two populations reached steady state and 90% of X_2_ cells were removed at Time 100. The orange lines are the total population size and the red and blue lines represent X_2_ fraction from the same models shown in *B*. The control model (blue and dashed orange lines) is the no-feedback model with compensated k_2_.

We simulated the model with an initial population containing X_1_ but not X_2_ with a parameter set adjusted such that the steady state ratio of the two populations is approximately 1:1 (Figure 3B, red), a ratio consistent with experimentally observed NE to non-NE transitions of SCLC cells^15^. We found that removing the feedback resulted in a lower steady state fraction of X_2_ due to the increased number of cells converting from X_2_ to X_1_ (Figure 3B, black). While this might suggest a role of the feedback signaling (Figure 3A, red) in facilitating the formation of mesenchymal-like X_2_ cell population, one can argue that this performance advantage can be simply achieved by adjusting the basal conversion rate constant *k*_2_ or other parameters. We therefore tested another feedback-free model in which there was a compensatory reduction of *k*_2_ (Figure 3B, blue) that produced the same 1:1 steady state ratio for the two types of cells. We found that the model with the inter-subtype signaling needed a significantly shorter time to achieve the equilibrium of the two subpopulations compared to the perturbed model that achieved the same level of heterogeneity (approximately three days shorter than control. See Methods.). We found that this acceleration was consistent in a range of steady state (target) ratios of the two subpopulations, but it was more prominent when the two subpopulations were comparable in size (Figure 3C). In addition, with the same range of the scanned k_2_ values, the model with the feedback had a narrower range of steady state compositions of the two populations (Figure 3C). This suggests that the inter-subtype feedback can be helpful to support desired phenotypic compositions in facing variations of kinetic rate constants. Interestingly, both acceleration and stabilization properties of the feedback were observed when we simulated a heterogeneity restoration from a purely M-like population that reflects a biologically plausible mesenchymal-to-epithelial transition (or a non-NE to NE transition in SCLC) (Figure 3D). Next, we asked whether cell-composition stabilization is a general behavior when all kinetic rate constants are perturbed simultaneously. We generated 1000 parameter sets, each with randomly chosen parameter values from a uniform distribution bounded by ± 50% deviations from the basal ones (Figure 3E). We found that the model with inter-subtype feedback had a small variation of phenotypic composition compared with the control model with respect to the global perturbation of parameters (control standard deviation (SD): 10.27%, SD with feedback: 5.52%) (Figure 3E, top). The feedback model also had shorter saturation time for heterogeneity restoration, as expected (*p*<10^−15^, Mann-Whitney U test) (Figure 3E, right).

We hypothesized that the acceleration function of the feedback loop may help the tumor cell population to re-establish a heterogeneous population under a subtype-specific “fractional condition,” i.e., a depletion of one of the subtypes. Indeed, after we perturbed a population already at equilibrium with equal numbers of each state (X_1_:X_2_ ratio of 1:1) by removing 90% of X_2_ cells from the system, the model with feedback recovered more rapidly compared to the feedback-free model (Figure 3F). Taken together, our mathematical model suggests two population-level functions of inter-subtype communication that we inferred from single-cell data: it accelerates the acquisition of heterogeneous tumor cell population from either a relatively homogeneous initial or out-of-equilibrium population, and it renders lower sensitivity of the steady state phenotypic compositions with respect to the perturbations of kinetic rate constants.

### SCLC utilizes autocrine and paracrine signaling to maintain epithelial state

We next asked how the cross-talk between the different states affects the epithelial state. In the RPM dataset, our analysis revealed that the epithelial state is sustained through a combination of paracrine and autocrine signaling mechanisms. We applied our analytical pipeline to construct a network for the Day 7 epithelial, SCLC-A2 subtype (Figure 4A). The network captured the activation of several epithelial marker genes. We observed the activation of the epithelial marker *Cdh1* in both the Day 7 and Day 11 network (Supplementary Figure 7). Additionally, the Day 7 network showed the activation of the SCLC-A2 marker, *Ascl1*. Notably, we identified *Sp1* as a key transcription factor involved in the activation of several epithelial markers. Interestingly, *Sp1* was also present in the SCLC-P/Y network, suggesting its potential to influence both NE and non-NE cell fate determination.

**Figure 4.**
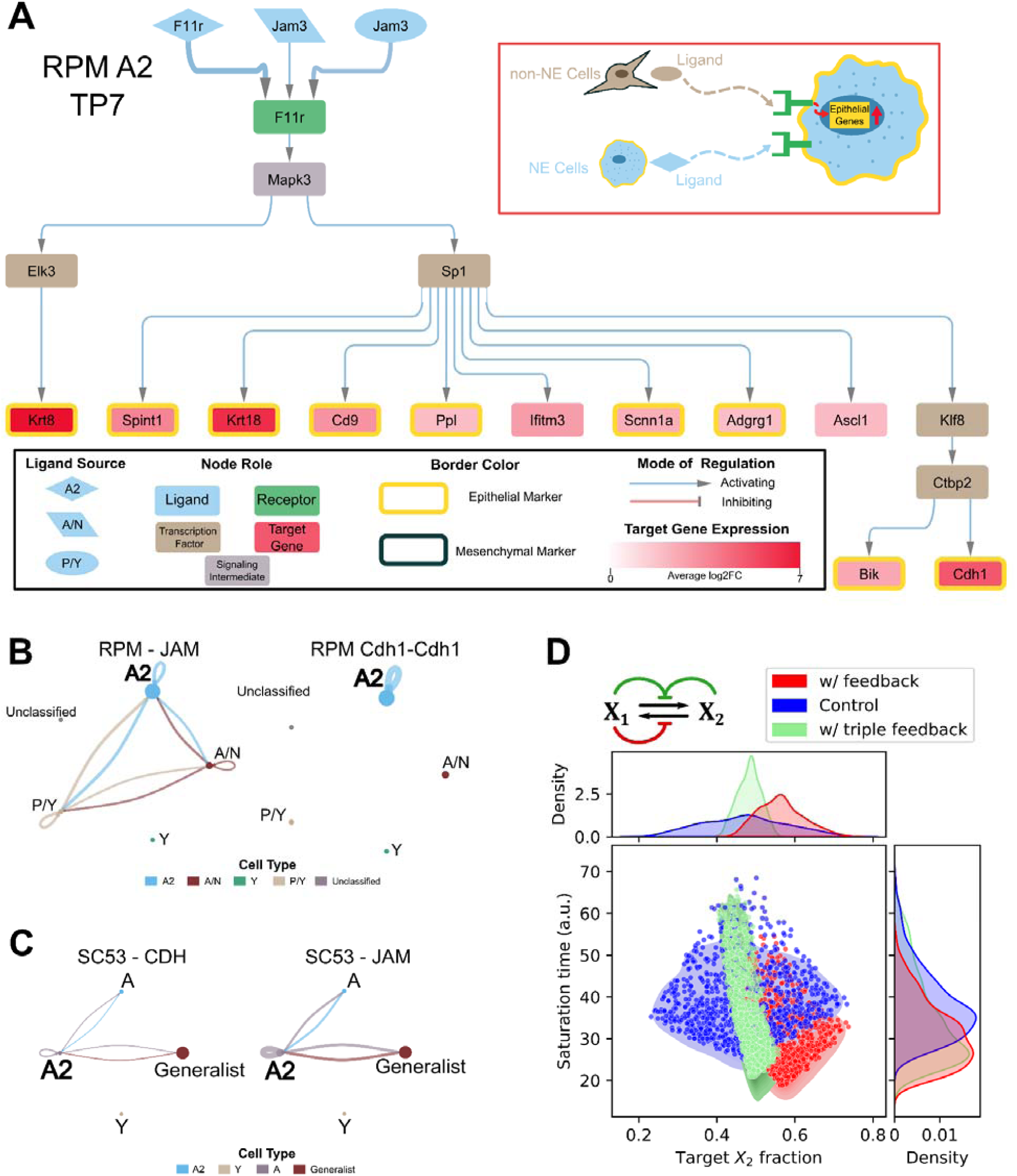
Epithelial Phenotype is Maintained by Autocrine and Paracrine Signaling. A) RPM TP7 SCLC-A2 cell type inferred signaling network. The inset indicated with a red rectangle is an illustrative summary of the network. The legend of the network is shown in the inset black rectangle. B) From the RPM data, the inferred JAM signaling pathway is shown on the left and the inferred L-R interaction of CDH1 is shown on the right. The Cell Type legend on the bottom contains the cell type each dot represents. C) Inferred CDH signaling pathway (left) and JAM pathway (right) from the human SCLC SC53 dataset. D) Performance of a model with 3 types of feedback (red and green arrows) in terms of the saturation time and phenotypic composition (measured as fractions of X_2_ cells). Performance of the triple-feedback model is shown in blue. A single-feedback model containing the E-to-M signal only (red) and a feedback-free model (blue) were used for comparison. 1000 random parameter sets were used for each model (see Methods for basal parameter values and ranges). Initial conditions: X_1_ =10, and X_2_=0.

Similar to observations in the SCLC-P/Y network, the SCLC-A2 network revealed the participation of both NE and non-NE cells in maintaining the epithelial state within SCLC-A2 cells. This interplay is also evident in the other inferred signaling pathways (Figure 4B). Specifically, the inferred interaction involving *Jam3* ligands within the JAM signaling pathway is consistent with prior work demonstrating *Jam3*’s role in establishing the epithelial phenotype^60^. Furthermore, we identified the CDH1 pathway as being activated by autocrine signaling within SCLC-A2 cells. The presence of CDH1 and JAM signaling pathways was also observed in human SCLC SC53 samples, operating in a manner similar to our findings in the SCLC-A2 network (Figure 4C and Supplementary Figure 8). This suggests that the maintenance of the SCLC epithelial state involves signaling interactions between epithelial and mesenchymal cells.

When applying the signaling network pipeline to the epithelial states of the other cancers, a network could only be generated for the colon cancer epithelial state but not the HER2 epithelial states (Supplementary Figure 6). The colon cancer network does not capture the activation of any of the overexpressed epithelial genes present within this cell type. For the HER2 epithelial states, a causal inference network could not be generated from the receptors to transcription factors. However, the less comprehensive nature of the colon cancer network and the inability to generate a network for the HER2 epithelial state could be due to additional mechanisms playing a role in either maintaining or destabilizing the epithelial state. This highlights the different levels of influence that cell–cell communication may play in maintaining the intratumoral heterogeneity, with the epithelial SCLC state being more sensitive to these signals. Nonetheless, we examined the potential roles of the SCLC-specific autocrine and paracrine signaling at the cell population level with mathematical modeling. We built a triple-feedback model that incorporates three types of cell communications (Figure 4D, top). With 1000 sets of randomly generated parameter sets each of the triple-feedback and benchmark models (i.e. the feedback-free model and the single-feedback model described in Figure 3), we found that signals that the NE cells (E-like cells) received produced the narrowest distribution of the phenotypic composition among the three models when parameters were perturbed globally (SD with single-feedback: 5.52%; SD with triple-feedback: 2.83%) (Figure 4D, top). This indicates that these forms of communications can further enhance the robustness of the phenotypic composition. However, the feedback on NE cells dampened the acceleration of the re-equilibrium compared to the single-feedback model (*p*<10^−13^, Mann-Whitney U test) (Figure 4D. right). This suggests that the system has a tradeoff between the speed and the accuracy (subpopulation fractions) of (re-)establishment of cellular heterogeneity. Overall, we found that all types of cell communication-based feedback inferred from single-cell data had pronounced effects on the equilibrium of the heterogeneous cancer cell populations.

## Discussion

Elucidating the dynamics and mechanisms that govern phenotypic plasticity within cancer tumors is essential for developing therapeutic strategies and tackling two major unresolved clinical challenges: cancer metastasis and therapeutic resistance^10,11^. While considerable progress has been made in characterizing phenotypic plasticity at the gene expression level^12,58,59,61^, many aspects remain poorly understood. Identifying the mechanisms that drive intratumoral heterogeneity and regulate phenotypic plasticity is a critical step for effective cancer treatment, as different cell types within a tumor can respond differently to therapies^22,62–66^. In this study, we investigated whether intercellular communications play a role in controlling cell fate transitions and whether these interactions stabilize or destabilize cellular phenotypes. We applied a multiscale inference-based approach to different solid tumor scRNA-seq datasets to investigate the crosstalk between cell states and how they influence one another. Within SCLC, we found the pivotal role of intercellular signaling in maintaining the phenotypic diversity among the cancer cell population, particularly in the context of EMT. The inferred SCLC-P/Y signaling network captures the activation of *Myc and* Notch signaling, which is consistent with recent observations as the activation of these two components is seen within the non-NE subtypes^16,21^. Additionally, the networks capture the mesenchymal nature of SCLC-P/Y subtype^24^, as the activation of many differentially overexpressed mesenchymal markers are present within the network. Furthermore, this mesenchymal phenotype is influenced by both mesenchymal P/Y cells and the more epithelial SCLC-A2 cells through the activation of EMT pathways Notch and Jam. The SCLC-A2 network displays similar patterns in that it utilizes both paracrine and autocrine EMT signals to maintain its epithelial state. Among several cancer types that we analyzed, this epithelial state maintenance mechanism appears to be unique to SCLC and specific to the SCLC-A2 subtype.

Applying this multiscale methodology to the colon and HER2 cancer datasets yielded less-comprehensive networks. One possible explanation for this difference is that the tumor microenvironment (apart from tumor cell heterogeneity) may play a larger role in influencing phenotypic plasticity in colon and HER2 breast cancer. In all three datasets, we only examined the intercellular communications within the cancerous population. In SCLC, the cell–cell communication between the cancerous cells appears sufficient enough to influence the cellular phenotypes. However, this does not appear to be the case with colon and HER2 cancers, where other factors in the tumor microenvironment may have a greater impact on tumor cell plasticity^67,68^. The HER2 data was derived from an *in vivo* setting and this context introduces greater complexity, as interactions with surrounding non-cancerous cells could significantly shape the phenotypic diversity observed. Alternatively, technical differences in how tumor cells are classified could affect differences in inferred networks. Our method requires sufficient cellular diversity (i.e. number of cells for each cell state or significant differences in gene expression between cell states) within the dataset in order to make inferences. The cancer cells in SCLC have been categorized via archetype analysis, which identifies the most relevant features to distinguish the cell classes in high-dimensional feature space. The cells in the colon cancer dataset were FACs sorted based on EpCam expression and they were assigned epithelial and mesenchymal scores through gene set enrichment analysis. The HER2 cell identities were inferred using single-cell trajectory analysis in which the terminal branches were used to annotate cells based on the differential expression of known marker genes. These differences in classification methods could influence the resolution of cell states and the inferred diversity within the datasets, potentially limiting the ability to detect nuanced intercellular communication networks in colon and HER2.

Phenotypic diversity in cell populations can be supported by both autonomous and non-autonomous mechanisms. Autonomous mechanisms include intrinsic transcriptional fluctuations, which can stochastically initiate phenotypic transitions^58^, as well as multi-stable regulatory networks, where cells can switch between stable phenotypic states based on underlying network architectures^69^. Additionally, post-transcriptional mechanisms provide another layer of intrinsic regulation^70^. However, while these autonomous processes can trigger phenotypic transition, it may be difficult to maintain this phenotypic diversity over time. Therefore, non-autonomous mechanisms may be required, such as paracrine/autocrine signaling which can play a critical role in reinforcing phenotypic heterogeneity through intercellular communication^71–73^. Our findings highlight the importance of these non-autonomous signaling mechanisms, demonstrating that cell–cell interactions can be essential for sustaining the intratumoral phenotypic heterogeneity.

Cell–cell interactions are fundamental to shaping and maintaining multicellular structures and tissue integrity. Among these, cell–cell communications play a pivotal role in coordinating cellular behavior across short and long distances within tissues. Here, we inferred cell–cell communications from scRNA-seq data, and while cell–cell communication methods based on scRNA-seq can predict short-range communications^68,74^, they are limited by the lack of spatial context and can introduce significant false-positives^75^. Integrating spatial transcriptomics with scRNA-seq offers a promising avenue to overcome these limitations, as it preserves the spatial arrangement of cells, allowing for more accurate inference of both short- and long-range communications^76,77^. Future work will benefit from leveraging these integrative approaches as it will refine our understanding of the role of cell–cell communications in maintaining intratumoral heterogeneity within tumors. In addition, this work focused on signaling with ligand-receptor interactions, which can in turn contribute to nonhomogeneous spatial patterns at the tissue scale. It will be of interest to study the reciprocity of cell-cell communications and tissue patterning in future modeling work.

Intercellular signaling is known to contribute to the intratumoral heterogeneity within SCLC, with the activation of Notch signaling resulting in NE to non-NE cell fate switching in 10–50% of tumor cells^15^. Additionally, the non-NE subtype exhibits a reduced proliferative rate but relatively greater chemoresistance, and these cells support the growth and survival of the NE subtype within admixed tumors^18^. This dynamic interplay between NE and non-NE subtypes highlights the role of intercellular signaling in maintaining a functional heterogeneity that benefits tumor survival and progression. A recent study suggests that SCLC subtypes not only coexist but may actively cooperate to optimize essential tumor functions, with NE and non-NE cells interacting in mutually beneficial ways to foster tumor growth and adapt to changing external conditions, such as treatment^21^. Additionally, it has been suggested that non-genetic mechanisms, such as cell-cell interactions between SCLC cell types, provides the capability for some tumors to reemerge once therapy is withdrawn through commensal niche-like interactions, where one cell type fosters the growth or survival of another^22^. These cooperative interactions are critical for maintaining the phenotypic diversity needed for tumor adaptability. Importantly, the rate at which phenotypic heterogeneity reaches equilibrium is likely driven by such cooperative mechanisms, enabling tumors to rapidly adapt by leveraging the distinct but complementary functions of different cell types. Disrupting these signaling networks or undermining the cooperative interactions between subtypes could impair the tumor’s ability to maintain this adaptability. thereby enhancing therapeutic efficacy.

We are only beginning to uncover the role in cancer of non-cell-autonomous signaling on phenotypic plasticity—a key driver of tumor progression and therapeutic resistance. Our results indicate that SCLC tumor cells have significant responses to extracellular signals emanating from different tumor cell subtypes and that transitions to mesenchymal phenotypes (especially SCLC-P/Y) are enhanced by ligands released from both NE and non-NE sources. Additionally, and somewhat surprisingly, the more epithelial SCLC-A2 state is likewise stabilized by signals from both NE and non-NE sources. Our mathematical model of the dynamics of cell state heterogeneity equilibration suggests that feedback mechanisms dependent on intercellular signaling are important modulators of how quickly and how accurately equilibrium can be reestablished. Altogether, our results support an important role for intercellular communication in controlling the dynamics of establishing equilibrium of intrinsic tumor cell state heterogeneity.

## Conclusions

Single cell-based inference approach reveals a key role of intercellular signaling in maintaining intratumoral heterogeneity. Cell-cell communications reinforce cell state transitions in multiple cancers at the single-cell level, and they support phenotypic heterogeneity at the population level by accelerating re-equilibrium and conferring robustness of population compositions.

## Methods

### Single-cell RNA-Sequencing Data

Single-cell RNA sequencing data were downloaded from Gene Expression Omnibus (GEO) at GSE149180 (RPM mouse tumor time course)^16^, GSE138474 (Human SCLC CDX)^25^, GSE154930 (HCT116 colon cancer)^34^, and GSE152422 (HER2 breast cancer mouse isoform). RPM mouse tumor dataset was preprocessed as described by Groves et al.^21^ Python package *Scanpy* (version 1.8.0) was used for filtering and normalization of total counts. Log-transformation was performed using the *log1p* function from the *Numpy* (version 1.17) package and scaling was done using *Scanpy*.

Human CDX data were preprocessed as described by Gay et al.^78^ Cells were filtered to remove non-tumor cells. Only the SC53 tumors were used in this analysis. *Scanpy* was used to normalize the total counts by cell and the data was then log-transformed.

Cell type annotation for RPM and SC53 datasets was performed as described by Groves, et al.^21^ Briefly, archetype analysis was applied to gene expression data, allowing for flexible characterization of the transcriptional landscape based on functional phenotypic features. This method approximates the cell phenotype space as a low-dimensional polytope that encapsulates the gene expression data. The vertices of the multi-dimensional space represent archetypes, each corresponding to a subtype optimized for a specific functional task. Cells are assigned to an archetype based on an archetype/optimization score, reflecting how well they align with that functional state. In the SCLC RPM dataset, some subtypes labels represent hybrid states (SCLC-A/N and SCLC-P/Y), indicating cells that exhibit transcriptional characteristics of two neighboring archetypes. There are also Unclassified cells present in this dataset, which are cells that are not strongly associated with any of the defined archetypes. In our study, we were able to assign more Unclassified cells into one of the archetypes by lowering the archetype assignment score threshold. Additionally, within the SC53 dataset, there are Generalist cells which are a non-specific cell type from archetype analysis and they do not fall into one of the archetype vertices. Instead, they occupy a more central position within the archetype space.

The deposited colon cancer data was already preprocessed as described by Sacchetti, et al.^34^ EpCAM^high^ and EpCAM^low^ raw count matrices were merged together in R and processed for downstream analysis using the *Seurat* package. Dimension reduction was performed using PCA, tSNE, and UMAP. Cell type annotation was performed based on EpCAM expression. Cells were also assigned epithelial and mesenchymal scores which were computed using gene set enrichment scoring packages.

HER2 breast cancer data was preprocessed as described by Ginzel, et al.^35^ Raw count matrices were processed using *Seurat* (version 4.0.0). Scores for S-phase and G_2_-M cell-cycle using the *CellCycleScoring* function. Data were log-normalized and scaled after regressing out total UMI counts, percent mitochondrial gene expression, and cell-cycle phase. Dimension reduction was done using PCA and UMAP. Individual clusters were annotated based on known marker gene expression. Trajectory analysis was performed with *Monocle 2* (version 2.12.0) on a subset of the data which contains only the epithelial compartment, and cellular identities were inferred in the terminal branches using a list of curated gene sets obtained from literature search and gene enrichment analysis^35^.

### Epithelial-Mesenchymal Enrichment Analysis

Two methods were used to compute epithelial and mesenchymal scores: non-negative principal component analysis (nnPCA)^24,79^ and single-sample gene set enrichment analysis (ssGSEA)^80^. An EMT gene set was used for the enrichment analysis^33^. Since some of the datasets originated from mouse models, human gene symbols from the gene set were converted to their mouse orthologs using the function *convert_human_to_mouse_symbols* from *nichenetr*^81^.

### Connecting cell-cell communication to TFs and Downstream Target Genes

#### 1. Cell-Cell Communication Inference

For our pipeline, the *CellChat* (version 1.6.1) package in R was used to infer intercellular communications. *CellChat* can quantitatively infer and analyze intercellular communication networks from scRNA-seq data. The interactions were identified and quantified based on the differentially over-expressed ligands and receptors for each cell group and a mass action-based model is used to integrate all known molecular interactions, including the core interaction between ligands and receptors with multi-subunit structure, as well as any additional modulation by cofactors. For each cell type, we took every inferred L-R interaction in which that cell type is receiving a signal and used those interactions as one of the inputs for the signaling network inference in later steps.

#### 2. Differential Expression Analysis and Transcription Factor Activity Inference

The *CORNETO* signaling network inference algorithm requires TFs and an activity score as another input so we performed differential expression analysis and TF activity inference. Differential expression analysis was performed using *Seurat’s FindMarkers* function. The function was applied to each cell type within the dataset. The differential expression results were then filtered for significance (p < 0.05). The statistically significant differentially expressed genes were then used for downstream processes.

Transcription factor activity inference was performed using *decoupleR*’s (version 2.6.0) univariate linear model method. This method requires gene expression values and a gene regulatory network as inputs. *decoupleR* provides the *CollecTRI* network which comprises a curated collection of signed TF-target gene interactions, weighted based on the mode of regulation^82^. This method will fit a linear model for each cell and each TF in the network, predicting the observed gene expression based on the TF-target gene interaction weights^83^. In our pipeline, we use the average log fold-change values to infer the activity of the TFs. This allows us to compare differences in TF activity between the cell types. We utilized the CollecTRI gene regulatory network for the network input. This was done for each cell type, so the differential expression results used in the function were specific to that cell type. Once the model is fitted, the obtained t-value of the slope is the score. The list of inferred active TFs was filtered for significance (p < 0.05) and activity (score > 0). The resulting inferred active TFs serve as inputs for the next step in the pipeline.

#### 3. Signaling Network Inference

The LIANA+ framework (version 1.0.2) was utilized to infer the signaling network. The *find_causalnet* function in the *CORNETO* package (v0.9.1-a5) infers the signaling network from receptors to transcription factors. This will integrate prior knowledge of signed and directed protein-protein interactions from *OmniPath*^84^ to construct the smallest-sign consistent network that explains the measured inputs and outputs, a network inference problem originally formulated in *CARNIVAL*^85^.

*CORNETO* requires a prior knowledge graph, receptor input scores, TF output scores, and node weights. This incorporation ensures the resulting network captures both dataset-specific information and established biological knowledge. For the receptor input scores, we utilized the probability score inferred from *CellChat*. Given that some receptors were inferred to be involved in multiple L-R interactions, we added the probability scores for a receptor if the inferred signaling pathway was the same. All inferred receptors were used as inputs for the signaling network algorithm. For the TF output scores, we used only the inferred active transcription factors.

*CORNETO* requires a prior knowledge graph of protein-protein interactions in order to connect the receptors to the TFs. The prior knowledge graph was created using the LIANA+ function, *build_prior_network.* Intracellular signaling Interactions were obtained from OmniPath using the *OmnipathR* (version 3.8.2) package. Interactions were obtained from the *omnipath, kinaseextra,* and *pathwayextra* datasets. The datasets were filtered based on curation effort (curation_effort > 1). The prior knowledge graph was created using the filtered interactions, input receptors, and output transcription factors. In our pipeline, we modified the node weights based on cell-type specific differential expression. Node weights had to be scaled from zero to one so the *MinMaxScaler* function from the Python library, *scikit-learn* (version 1.3.1), was used to scale the average log2 fold-change values. *CORNETO* requires a solver and an optimizer so we used the *CVXPY* (version 1.3.2) backend and *GUROBI* optimizer (*gurobipy* version 10.0.3) within the signaling network algorithm.

CORNETO will return the smallest sign-consistent network explaining the measured inputs and outputs, connecting the receptors to TFs. From this resultant network, we reincorporate ligands based on the L-R interactions from *CellChat*. In our pipeline, we also add downstream target genes to the network based on the consistency of mode of regulation from upstream TF and differential expression of target gene. For example, an activating edge towards a target gene necessitates a positive log fold-change value within the target gene in order for the gene to be added to the network. We used cell state-specific differential expression results for each network. TF-regulon interactions were obtained using *OmnipathR* via the *import_transcriptional_interactions* function. The interactions were filtered so that only SCLC^21^ (only for RPM and SC53 dataset) and EMT^33^ marker target genes were contained within the interaction list. Additionally, the list was filtered to contain experimentally validated interactions (curation_effort > 0).

The code of our new multiscale inference pipeline is available at https://github.com/DanielL543/scRNA_seq_multiscale_inference.

### Statistical Analysis of Networks

The cell-type specific differential expression results were used to determine the proportion of differentially expressed genes being in network vs out of network and whether the gene is lineage supporting or not. This was done using a 2×2 contingency table where the two variables are (1) the number of differentially expressed genes supporting or not supporting the cell lineage and (2) the number of differentially expressed genes in the network or out of the network. Lineage supporting genes were determined based on the previously reported phenotype of the cell type. Fisher’s exact test was then performed to determine the significance of a differentially expressed being in network and being lineage supporting (See Supplementary Table 1 for contingency tables of networks).

### Cell Line Authentication

This work does not involve experiments with cell lines. The authentication and check for contamination are therefore not required.

### Randomization

This work does not involve selection of subjects for experiments. The randomization procedure is therefore not required.

### Mathematical Modeling

For modeling the inter-subtype feedback that we identified as a conserved mechanism across cancer types, the ODE system shown in Eq 1 was used to simulate a cell population containing two subtypes of cancer cells. A representative parameter set was used for simulations: *r*_1_ = 0.01, *r*_2_ = 0.001, *k*_1_ = *k*_2_ = 0.04, *K*_2_ = 0.5, and *n*_2_ = 6. For feedback-free models (control), *K*_2_ was set to 100. *k*_2_ was adjusted to allow a feedback-free model (*k*_2_ compensated model) to achieve desired fraction of X_2_ cells. The model is dimensionless. One time unit in the model corresponds to 5 hours approximately. The initial conditions for the scenario of a purely E-like cell population are *X*_1_ = 10, *X*_2_ = 0, whereas initial conditions *X*_1_ = 0, *X*_2_ = 10 were used to simulate the opposite scenario. To examine the performance of the model with random parameter values, 1000 parameter sets were generated with a random sampling from uniform distributions bounded by (0.5, 1.5) where is the basal value for each parameter described above.

To incorporate additional feedback regulations that we inferred from the time course data for SCLC, we used the following ODEs to describe a “triple-feedback” model:

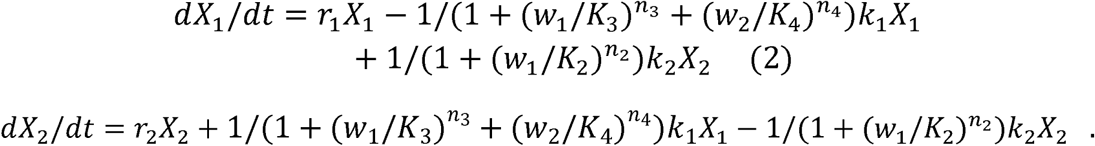

In addition to the variables and parameters in the single-feedback model (Eq 1), *w*_2_ is the fraction of *X*_2_ cells (*w*_2_ = *X*_2_/(*X*_1_ + *X*_2_)); *K*_3_ is the threshold of the autocrine effect on cell state transition from E-like cells to M-like cells; *K*_4_ is the threshold of the M-to-E signal effect on cell state transition from E-like cells to M-like cells; *n*_3_ and *n*_4_ describe the nonlinearity of the two effects, respectively. To randomly sample parameter values, basal values *K*_3_ = 0.9, *K*_4_ = 0.5 and *n*_3_ = 4 =6 were used with the same scheme for the single-feedback model.

## Ethics approval and consent to participate

This work does not involve human participants or animals.

## Consent for publication

This manuscript does not contain any individual person’s data.

## Availability of data and materials

The code for reproducing all results of this work is available at https://github.com/DanielL543/scRNA_seq_multiscale_inference. No new data is generated in this work.

## Authors’ contributions

Developed computational methods: D.L. and T.H.. Analyzed data: D.L., D.R.T. and T.H.. Wrote the manuscript: D.L., D.R.T. and T.H.. All authors read and approved the final manuscript.

## Acknowledgements

The authors thank the members of Tian Hong’s laboratory for helpful comments.

