## Supplemental Figures for "Intercellular signaling reinforces single-cell level phenotypic transitions and facilitates robust re-equilibrium of heterogeneous cancer cell populations"

### Supplementary Figure 1

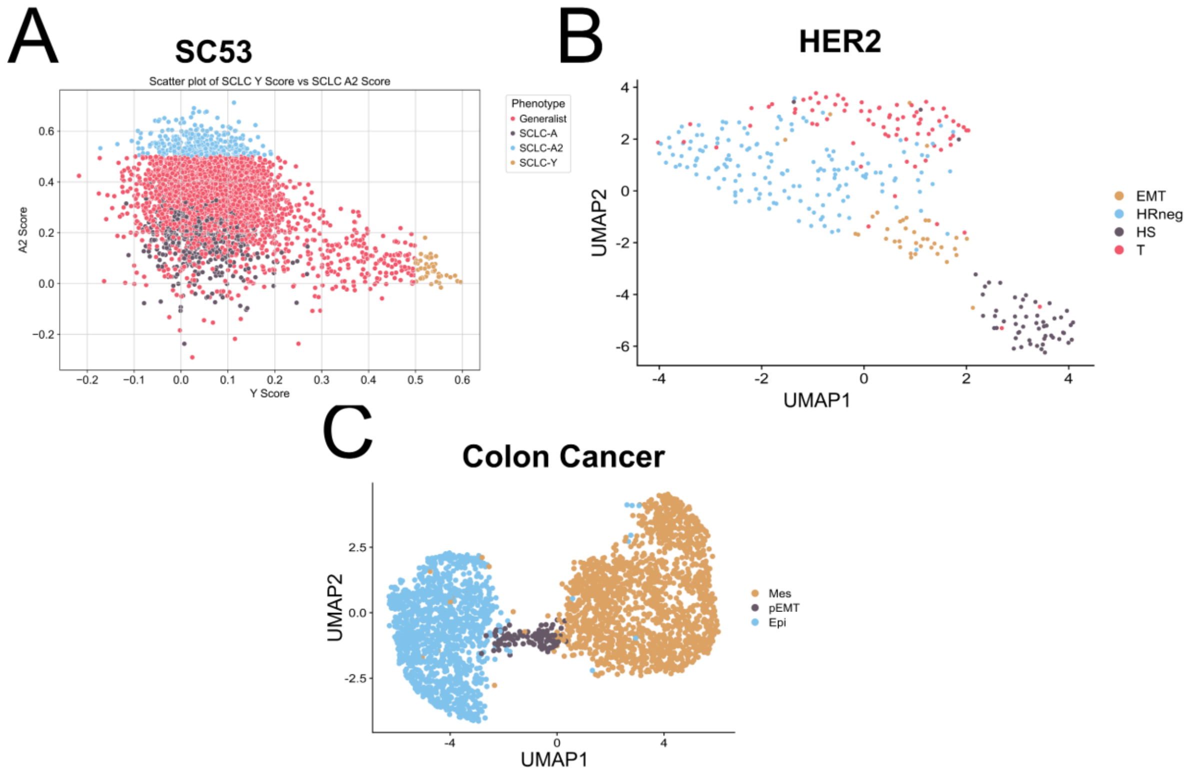


*Supplementary Figure 1:* ***Plots displaying the different cell types/cell-states present in SC53, HER2 and colon cancer data.*** *A) Cell types present in the CTX SC53 dataset. Cell types were determined by archetype analysis. Archetype analysis scores for the Y and A2 subtype are used to plot the different cells. B) cell types from the t039232 sample of the HER2 dataset. Cell types were determined based on the enriched expression of cell type markers from the terminal branches of trajectory analysis. C) Cell types from the colon cancer datasets (HCT1116 cell line). Cell types are characterized by EpCam status from FACS and scRNA-seq analysis*

### Supplementary Figure 2

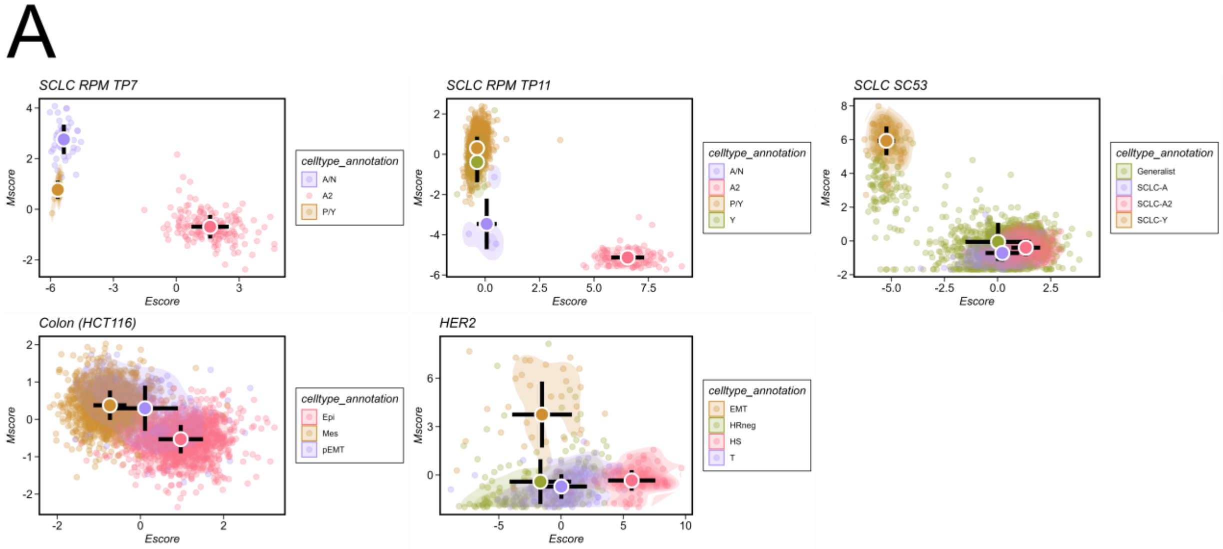


*Supplementary Figure 2:* ***Gene set enrichment comparisons of epithelial and mesenchymal populations across datasets.*** *A) Scatter plots of the results from gene set enrichment analysis using non-negative principal component analysis. Each dot represents a cell in that dataset. X-axis and y-axis represent enrichment scores of epithelial and mesenchymal genes, respectively.*

### Supplementary Figure 3

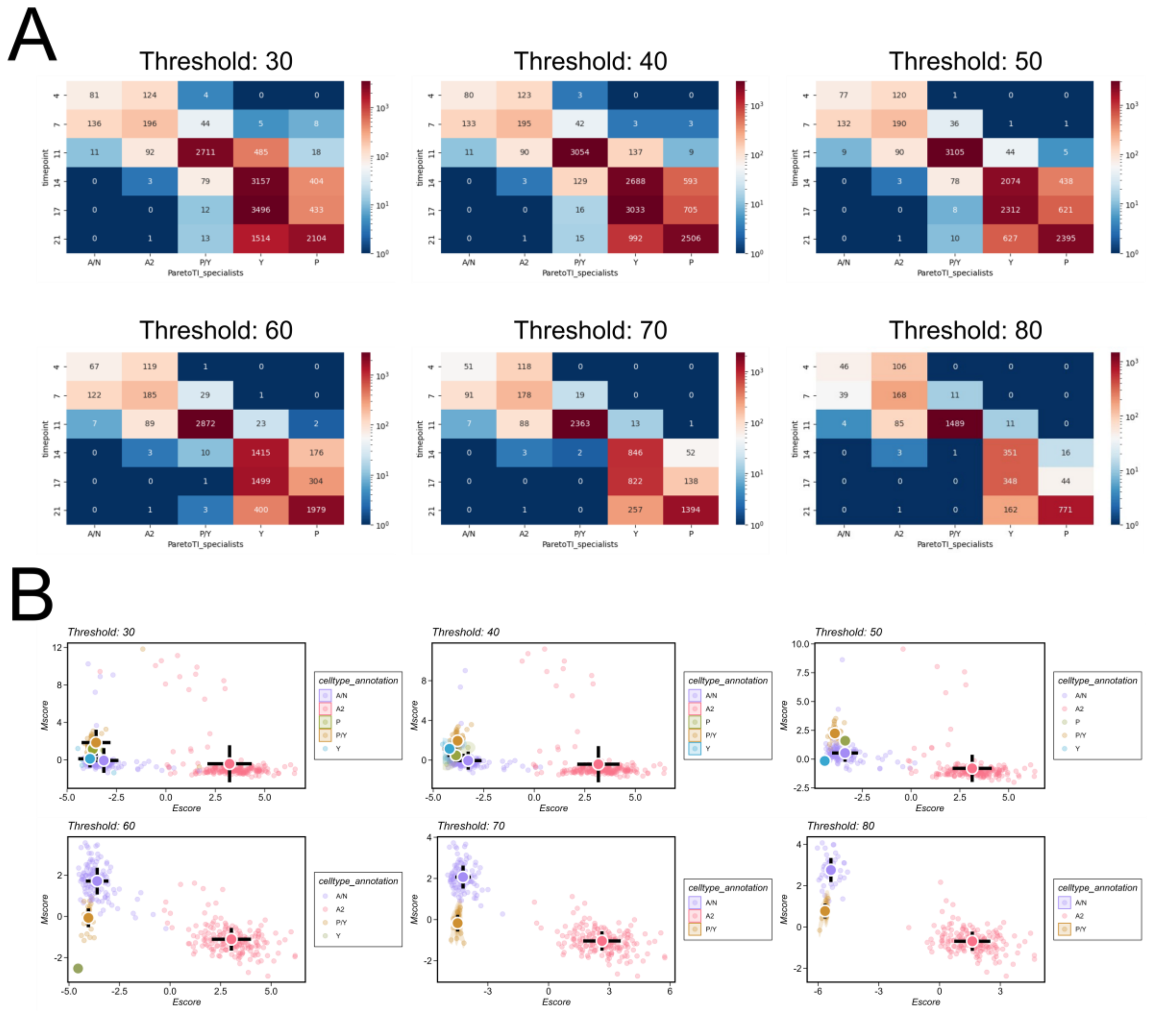


*Supplementary Figure 3:* ***Comparison of cell type annotations in SCLC RPM across different thresholds for archetype assignment****. A) Differences in cell counts across threshold values. X-axis represents the different types and timepoint is on the y-axis. The numeric value in each rectangle represents the number of cells for that given cell type at the timepoint. B) Scatter plots of nnPCA gene enrichment analysis at timepoint 7 across different archetype threshold values. Each dot in the plot represents a single cell. X- and y-axis represent enrichment score of epithelial and mesenchymal genes, respectively.*

### Supplementary Figure 4

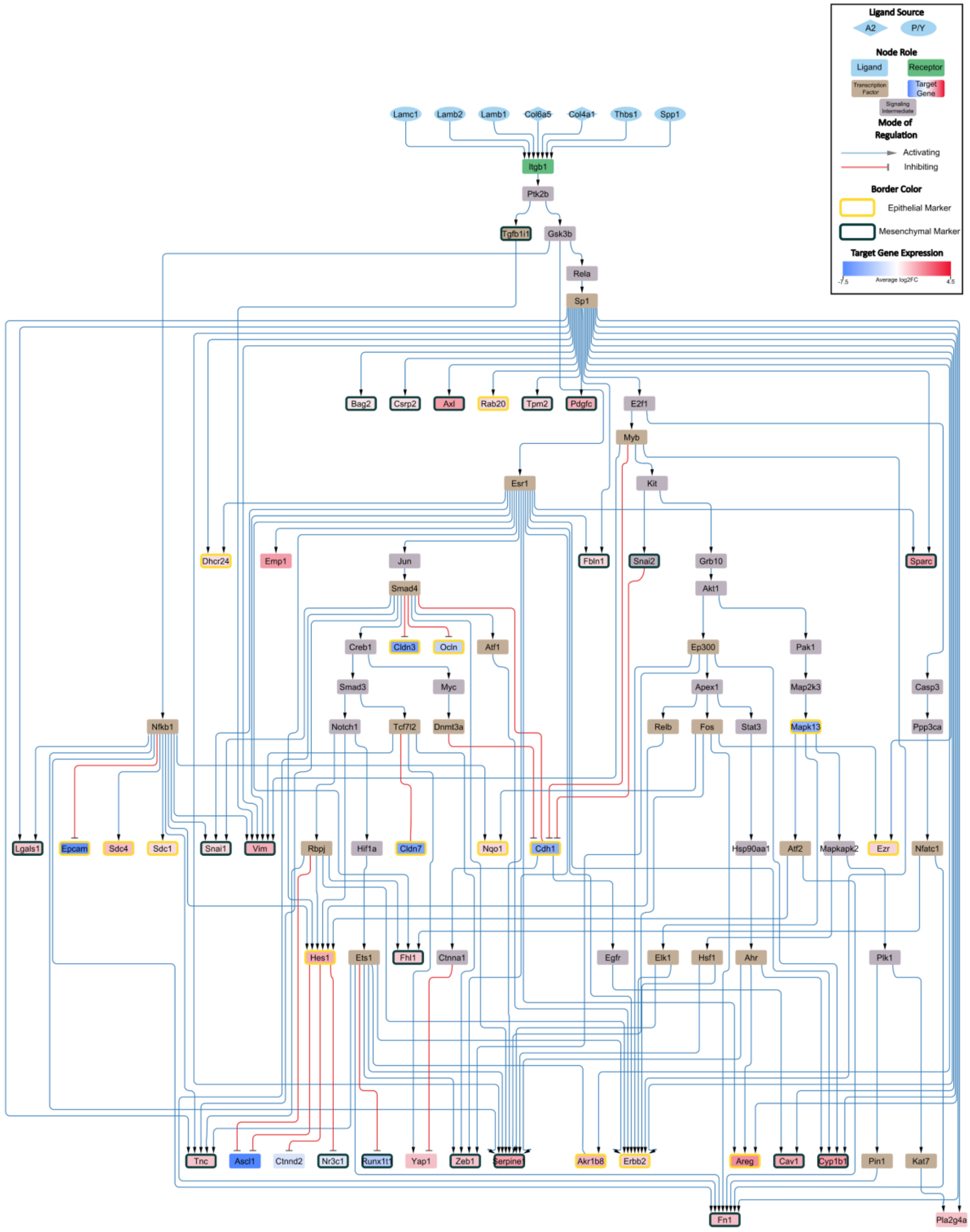


*Supplementary Figure 4: **P/Y cells at timepoint 11 utilize signals from NE and non-NE cell types to reinforce mesenchymal phenotype**. The node color represents the the role of the node/gene in the network, with the target genes colored based on log-fold change value from dark blue to red. Yellow borders around nodes represent epithelial markers and black borders are mesenchymal markers. Line width from ligands (light blue nodes) to receptors (green nodes) represent inferred interaction strength from CellChat.*

### Supplementary Figure 5

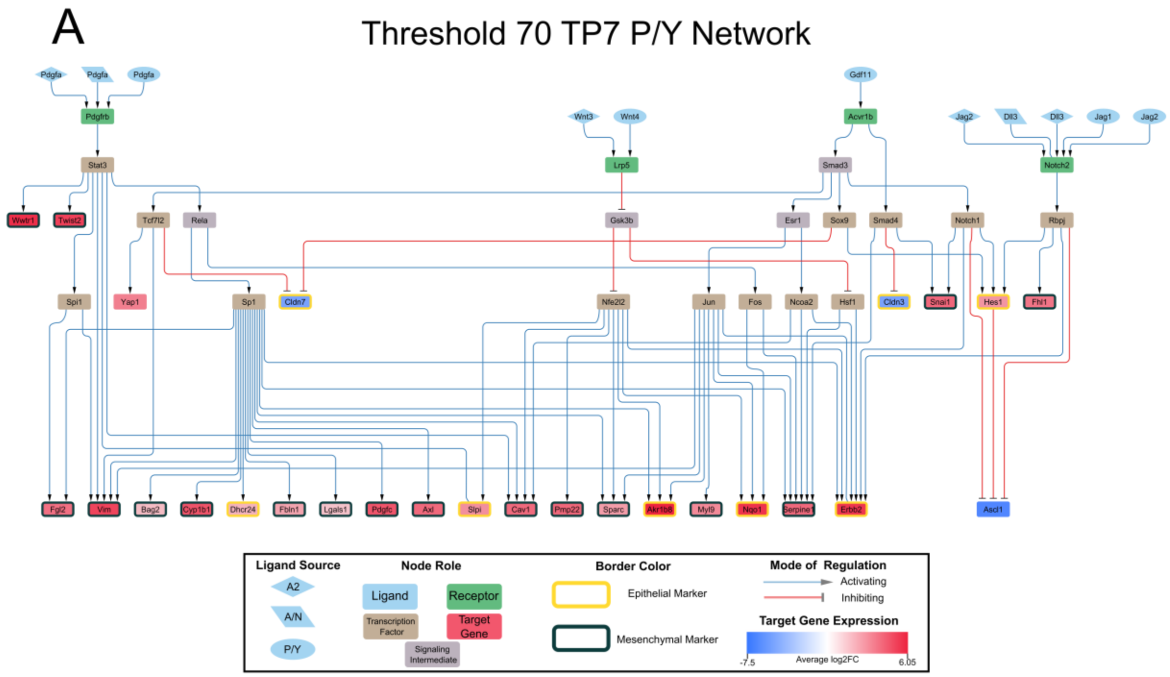


*Supplementary Figure 5:* ***Signaling network for the SCLC-P/Y subtype at TP7 when the archetype assignment threshold is lowered by 10****. Black box contains the legend for the figure. Light blue nodes are ligands, green nodes are receptors, grey nodes are signaling intermediates, tan nodes are TFs, and target genes are colored on a blue-red spectrum based on average log2FC. Blue edges and red edges represent an activating or inhibiting interaction, respectively. Ligand nodes have different shapes which represent different SCLC subtype sources.*

### Supplementary Figure 6

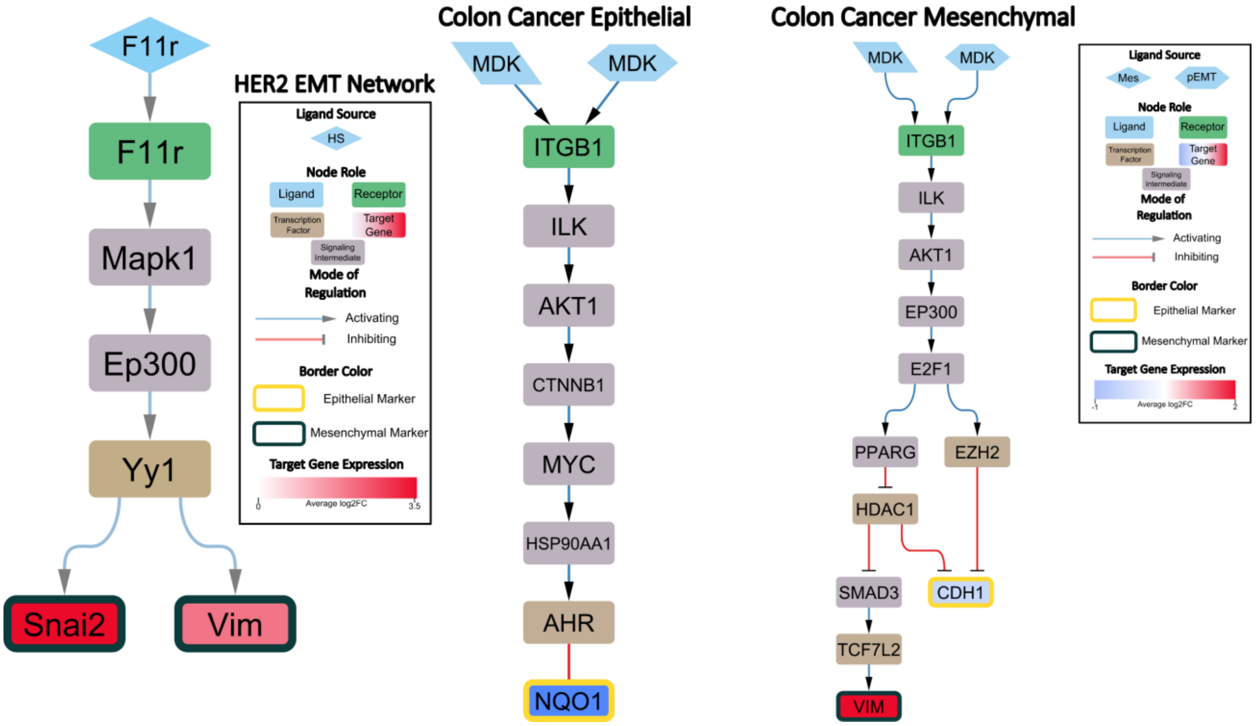


*Supplementary Figure 6:* ***Signaling networks created from multiscale inference approach for HER2 and colon cancer data.*** *A) The mesenchymal (EMT) cells in the HER2 dataset utilize signals from the more epithelial-like cells to maintain its phenotype. B) Signaling network for the epithelial cells in the colon cancer dataset. C) Inferred mesenchymal cell type signaling network in the colon cancer dataset.*

### Supplementary Figure 7

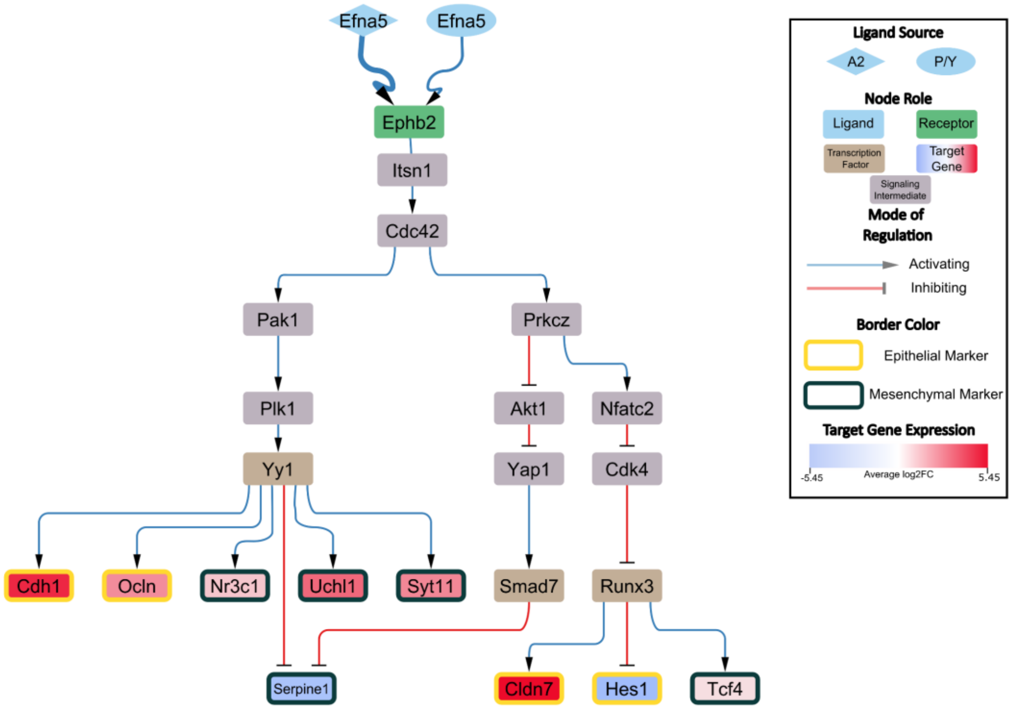


*Supplementary Figure 7:* ***The A2 cells in RPM TP11 data use signals from both NE and non-NE cells to maintain its epithelial phenotype.*** *Ligands are from A2 and P/Y cells.*

### Supplementary Figure 8

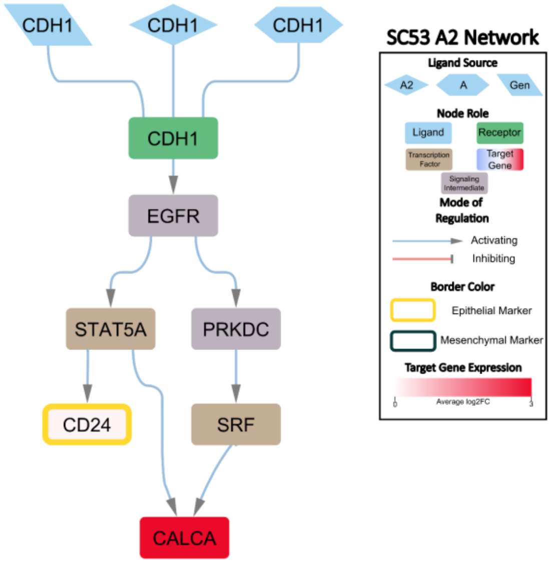


*Supplementary Figure 8:* ***Interactions between A2 cells help to reinforce the epithelial phenotype in SC53.*** *Ligands are from A2, A and Generalist cells. CALCA is an A2 marker*
